# Negative allosteric modulation of CB1 cannabinoid receptor signaling decreases intravenous morphine self-administration and relapse in mice

**DOI:** 10.1101/2024.01.16.575900

**Authors:** Idaira Oliva, Fezaan Kazi, Lucas N. Cantwell, Ganesh A. Thakur, Jonathon D. Crystal, Andrea G. Hohmann

## Abstract

The endocannabinoid system interacts with the reward system to modulate responsiveness to natural reinforcers, as well as drugs of abuse. Previous preclinical studies suggested that direct blockade of CB1 cannabinoid receptors (CB1R) could be leveraged as a potential pharmacological approach to treat substance use disorder, but this strategy failed during clinical trials due to severe psychiatric side effects. Alternative strategies have emerged to circumvent the side effects of direct CB1 binding through the development of allosteric modulators. We hypothesized that pharmacological inhibition of CB1R signaling through negative allosteric modulation (NAM) would reduce the reinforcing properties of morphine and decrease opioid addictive behaviors. By employing i.v. self-administration in mice, we studied the effects of the CB1-biased NAM GAT358 on morphine intake, relapse-like behavior, and motivation to work for morphine infusions. Our data revealed that GAT358 reduced morphine infusion intake during the maintenance phase of morphine self-administration under fixed ratio 1 schedule of reinforcement. GAT358 decreased morphine-seeking behavior after forced abstinence. Moreover, GAT358 dose-dependently decreased the motivation to obtain morphine infusions in a progressive ratio schedule of reinforcement. Strikingly, GAT358 did not affect the motivation to work for food rewards in an identical progressive ratio task, suggesting that the effect of GAT358 in decreasing opioid self-administration is reward specific. Furthermore, GAT58 did not produce motor ataxia in the rota-rod test. Our results suggest that CB1R NAMs reduced the reinforcing properties of morphine and could represent a viable therapeutic route to safely decrease opioid-addicted behaviors.

## 1. Introduction

Opioid use disorder (OUD) remains at epidemic levels, affecting 16 million people and causing over 120,000 deaths annually from opioid overdose worldwide ^1^. Current medications for OUD that target the µ-opioid receptor, such as methadone or buprenorphine, face numerous challenges, including requirements of certified medical personnel, licenses, prior authorizations, and logistical hurdles ^2^. Additionally, these treatments are not risk-free or successful in all patients.

µ-opioid receptors and cannabinoid 1 receptors (CB1R) are G-protein coupled receptors (GPCRs) that are distributed in many of the same areas in the brain, including the classical reward system ^3^. Bidirectional interactions between these two GPCRs have been reported on reward, tolerance, and other addiction-related behaviors ^4,5^. These observations have prompted a growing interest in targeting the endocannabinoid system as a potential avenue for developing alternative and/or adjunct treatments in OUD. Preclinical studies suggested that CB1R antagonists rimonabant (SR141716A) and AM251 were efficacious in reducing consumption of several drugs of abuse in rodents, including psychostimulants, opioids, alcohol, and nicotine ^5^. However, when rimonabant was used as an anti-obesity drug in people, adverse psychiatric side effects led to its withdrawal from the market and the termination of clinical programs on CB1 antagonists/inverse agonists. Since then, alternative strategies have emerged to circumvent the negative outcomes of direct CB1R binding through the development of allosteric modulators. Currently, very few preclinical studies have examined the therapeutic effects of CB1R negative allosteric modulators (NAMs) in drug addiction. The two most studied allosteric modulators of CB1R receptors are Org27569 and PSNCBAM-1. Org27569 attenuated cue- and drug-induced reinstatement of cocaine- and methamphetamine-seeking behavior in rats ^6^. PSNCBAM-1 attenuated the reinstatement of extinguished cocaine-seeking behavior in rats ^7^. However, questions have been raised regarding their selectivity given that, like rimonabant, Org27569 and PSNCBAM-1 caused anhedonia, decreasing food intake ^8,9^. Furthermore, both Org27569 and PSNCBAM-1 displayed a contradictory pharmacological profile, acting as an inverse agonist while also acting as a negative allosteric modulator of CB1R depending on the orthosteric agonists active in a given circuit ^10–12^.

We recently developed a novel CB1R NAM, GAT358, using a focused structure-activity relationship study on PSNCBAM-1, which displays functional selectivity in the β-arrestin assay over the cAMP formation (biased NAM) ^13^. Our previous studies demonstrated that the CB1R NAM GAT358 suppressed morphine reward (i.e., conditioned place preference (CPP)) ^14^. However, no prior research has yet examined the effects of a CB1 NAM on opioid reinforcement using operant behavior paradigms (which involve drug, context, cues, and an action that is performed to obtain the drug i.e., i.v. morphine self-administration).

In the present work we investigated the effects of the CB1R NAM GAT358 on the reinforcing properties of morphine by employing i.v. drug self-administration in mice, the gold standard for studying drug addiction. Our results collectively suggest that the CB1R NAM GAT358 reduces the reinforcing properties of morphine. CB1 NAMs such as GAT358 may represent a viable therapeutic route to decrease opioid-addicted behaviors and relapse while circumventing the adverse side effects of CB1R orthosteric binding.

## 2. Materials and methods

### 2.1. Drugs

GAT358 was synthesized in the laboratory of Ganesh Thakur in the Department of Pharmaceutical Sciences at Northeastern University (by LNC). GAT358 was dissolved in a vehicle containing 20% dimethyl sulfoxide (DMSO) and 80% of ethanol:Alkamuls EL-620:saline in a 1:1:8 ratio and administered i.p. at 10, 20 or 30 mg/kg in a final volume of 5 ml/kg injection. Dose selection was based on our previous studies showing that 20 mg/kg (i.p.) decreased morphine-induced increases in dopamine efflux in the NAc, blocked morphine-induced CPP, and decreased the oral consumption of oxycodone ^14^. Morphine hydrochloride was obtained from the National Institute on Drug Abuse (NIDA), dissolved in a physiological saline solution, and administered at 300 µg/kg/infusion in the i.v. drug self-administration paradigm.

### 2.2. Animals

All procedures were approved by the Institutional Animal Care and Use Committee at the Indiana University Bloomington. Adult C57/BL male mice (Envigo, IN), 15 weeks old at the start of the study, were group-housed upon arrival, given ad libitum access to water/food, and maintained on a 12-hour reversed light/dark cycle. Mice were subjected to at least 48-hours of acclimation to the animal facilities prior to starting any experimental procedure. All mice were single-housed for the entire duration of the experiments.

### 2.3. Jugular catheter surgery

Surgeries were performed under isoflurane anesthesia (2–3%, 1.5 L/min O_2_). A back mount cannula (315BM-8-5UP, P1 technology, Roanoke, VA) was connected to a 4 cm micro-renathane tubing (0.025-inch outer diameter, 0.012-inch inner diameter, Braintree Scientific Inc.) and a silicon bead (kit-silicone low viscosity, WPI, Sasasota, FL) was placed approximately 10 mm from the tip ^15^. Briefly, tubing was passed subcutaneously from the back of the mice to the neck and surgical silk sutures were used to anchor the catheter to the jugular vein above and below the silicone bead. Incisions were closed using tissue adhesive (Vetbond, 3M). Mice were allowed to recover for 1 week following the catheter surgery and before starting the morphine self-administration sessions. Catheters were flushed daily (20 µl) with a heparinized saline solution (100 units/ml) before and after the drug self-administration sessions.

### 2.4. i.v. morphine self-administration

Animals were tested once daily in 120 min sessions drug self-administration sessions. Operant chambers (Med Associate Inc., Georgia) were equipped with grid floors, a house light, left and right nose poke ports and an infusion pump. A morphine concentration of 300 µg/kg/infusion was used for these experiments ^16^. Each trial started with the house light ON and the guillotine doors opened allowing access to the nose poke ports. An action on the active nose poke port resulted in the house light OFF and the delivery of one morphine infusion paired with the active nose poke light (i.e., discrete cue). Actions in the inactive nose poke port had no scheduled outcomes. Specifically, the infusion pump and the nose poke light turned ON for approximately 2 s and 3.6 µl of morphine solution (2.5 mg/ml) were infused based on the mouse’s body weight. Each infusion was followed by a 30 s timeout period where actions on both nose pokes ports did not have any consequence. It is important to highlight that in our self-administration experimental design mice have not been pre-trained to nose poke for food and the drug consumption in our experiments is truly volitional. In the maintenance experiment, mice were first allowed to self-administer morphine for 10 consecutive days on a FR1 schedule until completing the acquisition phase, ensuring that they distinguished between the reinforced and the non-reinforced nose poke. Then, mice continued in FR1 until they had stable behavior. The criterion for stable morphine intake was reached when the mice received at least 10 infusions per session and exhibited less than 15% variance in the number of infusions for three consecutive days ^17^. After meeting this criterion, the animals received counterbalanced injections of GAT358 (20 mg/kg i.p.) or vehicle (i.p.) 20 min before the self-administration session. Mice were allowed to recover baseline infusion intake between treatments. We analyzed the number of infusions and the number of active/inactive nose pokes, and within-subjects comparisons were made between the session with vehicle and GAT358 pretreatments. In the relapse test, a different group of mice first self-administered morphine for 15 days on a FR1 schedule. Animals that achieved at least 10 infusions per session at the end of the 15 days, then experienced 21 days of forced abstinence (i.e., withdrawal) in their home cages followed by a single cue-induced drug seeking session (i.e., relapse test) in the operant chambers. To minimize the experimenter-induced stress on the relapse test day, animals were handled daily during the forced abstinence period. Mice were treated with GAT358 (20 mg/kg i.p.) or vehicle (i.p.) 20 min prior to the relapse test session. Subsequently, the subjects were placed back into to the drug self-administration operant chambers with an identical configuration as before; however, active nose poking resulted in the delivery only of the discrete cue previously paired with morphine (i.e., nose poke light), but no morphine was infused. We analyzed active/inactive nose pokes and made comparisons between vehicle and GAT358 groups. In the progressive ratio experiment, a different cohort of mice was trained to self-administer morphine for 10 days in FR1, 5 days in FR2 and 5 days in FR3. Animals were required to get at least 10 infusions before progressing to the next FR. Next, mice were subjected to a progressive ratio (PR) schedule of reinforcement where they increasingly had to make more effort in the same session to get an infusion (i.e., PR values were 0, 1, 2, 4, 9, 12, 15, 20, 25, 32, 40, 50, 62, 77, 95, 118, 145, 178, 219, 268, 328, 402, 492, 603, 737, 901, 1102, 1347, 1647, 2012). Mice continued in PR until achieving a stable intake of morphine. As previously mentioned, a stable morphine infusion intake was defined as less than 15% in the number of infusions across three consecutive days. In the following PR sessions, mice were counterbalanced treated with GAT358 (10 mg/kg i.p.), GAT358 (20 mg/kg i.p.), GAT358 (30 mg/kg i.p.) or vehicle (i.p.) 20 min before testing. Animals were allowed to recover baseline levels of infusion intake in between treated sessions to avoid any carryover effects between doses. We analyzed the number of infusions, active/inactive nose pokes and break point (i.e., last ratio completed or limit amount of “work” that a subject is willing to perform to obtain a reinforcer), and within-subjects multiple comparisons were made between treatments.

### 2.5 Rota-rod test

A separate cohort of otherwise naïve mice was first subjected to 30 min of acclimation to the testing room and subsequently placed on the rota-rod. Animals underwent 2 training days and 1 testing day on the rota-rod apparatus (IITC Life Sciences Inc., Woodland Hills, CA). The rota-rod apparatus was set to start rotating at 4 RPM, end at 40 RPM and employed a 300 s cutoff. The goal for the training days for each mouse was to stay on the drum of the apparatus for at least 30 s in 3 consecutive trials with a maximum of 6 trials a day and 20 min interval between trials. On the testing day, to qualify for testing, each mouse ran a single trial and was required to stay in the drum for at least 30 s. Animals that did not meet this criterion were excluded from the experiment. Mice that qualified for testing, experienced 2 baseline trials (i.e., pre-injection), and then were randomly treated with GAT358 (10 mg/kg i.p.), GAT358 (20 mg/kg i.p.), GAT358 (30 mg/kg i.p.) or vehicle (i.p.) 20 min before being subjected to another 2 trials (i.e., post-injection).

### 2.6. Food self-administration

The VEH-treated animals previously subjected to the rota-rod test were subsequently food restricted (1.7-2 g regular chow per day) until they lost a 10% of body weight (approximately 1 week). These animals remained in food restriction for the entire duration of the experiment. For food self-administration, we employed identical operant chambers to those used for drug self-administration. These chambers were also equipped with two nose poke ports, one designated as an active and the other one as an inactive nose poke. Each trial started with the house light ON. Actions in the inactive nose poke produced no consequences whereas the required actions in the active nose poke turned OFF the house light and delivered a 14 g sucrose pellet paired with a nose poke light cue. The reward delivery was followed by a 30 s timeout, during which the house light remained OFF and actions in both nose pokes had no scheduled outcomes. Once the timeout period concluded, the house light turned ON again and a new trial started. Mice were first trained in FR1, FR2 and FR3 with the requirement to earn 60 or more pellets before passing to the next FR. Mice were then trained in a PR (i.e., same PR values as for drug self-administration) until the achieve a stable number of pellets earned. As before, a stable pellet intake was defined as less than 15% in the number of pellets earned across 3 consecutive days. In the following PR sessions, mice received, in a counterbalanced manner, GAT358 (10 mg/kg i.p.), GAT358 (20 mg/kg i.p.), GAT358 (30 mg/kg i.p.) or vehicle (VEH, i.p.) 20 min before testing. Animals were allowed to recover previous amounts of food pellet intake in between treated sessions to avoid any carryover effects between doses. We analyzed the number of pellets earned, active/inactive nose pokes, and breakpoints, performing within-subjects multiple comparisons between treatments.

### 2.7. Statistical analysis

Data were analyzed by One-way or Two-way Analysis of Variance (ANOVA) followed by Dunnett’s or Bonferroni post-hoc tests. The Geisser-Greenhouse correction was applied to all repeated factors to adjust the lack of sphericity in repeated measures ANOVA. Significant differences between two groups were assessed using a two-tailed paired t-test. Drug self-administration training sessions, in which mice engaged in biting the morphine delivery tubing, were excluded from the study. All statistical analyses were performed using GraphPad Prism (GraphPad Software, La Jolla, CA). Data are presented as mean ± SEM and *p*<0.05 was considered statistically significant.

## 3. Results

### 3.1. GAT358 reduces morphine intake during the maintenance phase of self-administration

We first evaluated the impact of GAT358 (20 mg/kg i.p.) on morphine intake during the maintenance phase of drug self-administration (**Figure 1**). Mice received 10 training sessions under FR1 morphine self-administration corresponding to the acquisition phase of morphine self-administration, they continued self-administering morphine under FR1 until the infusion intake was stable and then received a GAT358 (20 mg/kg i.p.) and vehicle (**Figure 1A**). During the acquisition phase, mice escalated the number of morphine infusions intake (**Figure 1B**) and notably distinguished the reinforced and non-reinforced nose pokes across sessions (**Figure 1C**). In the maintenance phase, the number of infusions and the number of active nose pokes decreased with GAT358 (20 mg/kg i.p.) but not with vehicle (VEH, i.p.) (**Figure 1D-E**). Specifically, GAT358 (20 mg/kg i.p.) reduced morphine infusion intake (paired t-test, two-tailed, p=0.0399; **Figure 1F**). Likewise, the total number of active nose pokes was decreased by GAT358 (20 mg/kg i.p.) (paired t-test, two-tailed, p=0.0098; **Figure 1G**). No differences were found in the total number of inactive nose pokes when administering GAT358 (20 mg/kg i.p.) or vehicle (i.p.) (paired t-test, two-tailed, p=0.3121; **Figure 1H**). The total quantity of morphine consumed was lower during the sessions treated with GAT358 (20 mg/kg i.p.) compared to vehicle treatment (paired t-test, two-tailed, p=0.0288; **Figure 1I**). Similarly, GAT 358 (20 mg/kg i.p.) reduced the number of cumulative infusions compared to vehicle (**Figure 1J**). Two-way ANOVA revealed a main effect of time (F_24,144_=25.7, p<0.0001), treatment (F_1,6_=6.641, p=0.0419) and the interaction was significant (F_24,144_=1.665, p=0.0357). Bonferroni’s post-hoc multiple comparisons indicated substantial differences in the number of infusions starting at 20 min bin (p=0.0162), 25 min bin (p=0.0015) and continuing from 30 to 120 min bins (p<0.0001) (**Figure 1J**).

**Figure 1.**
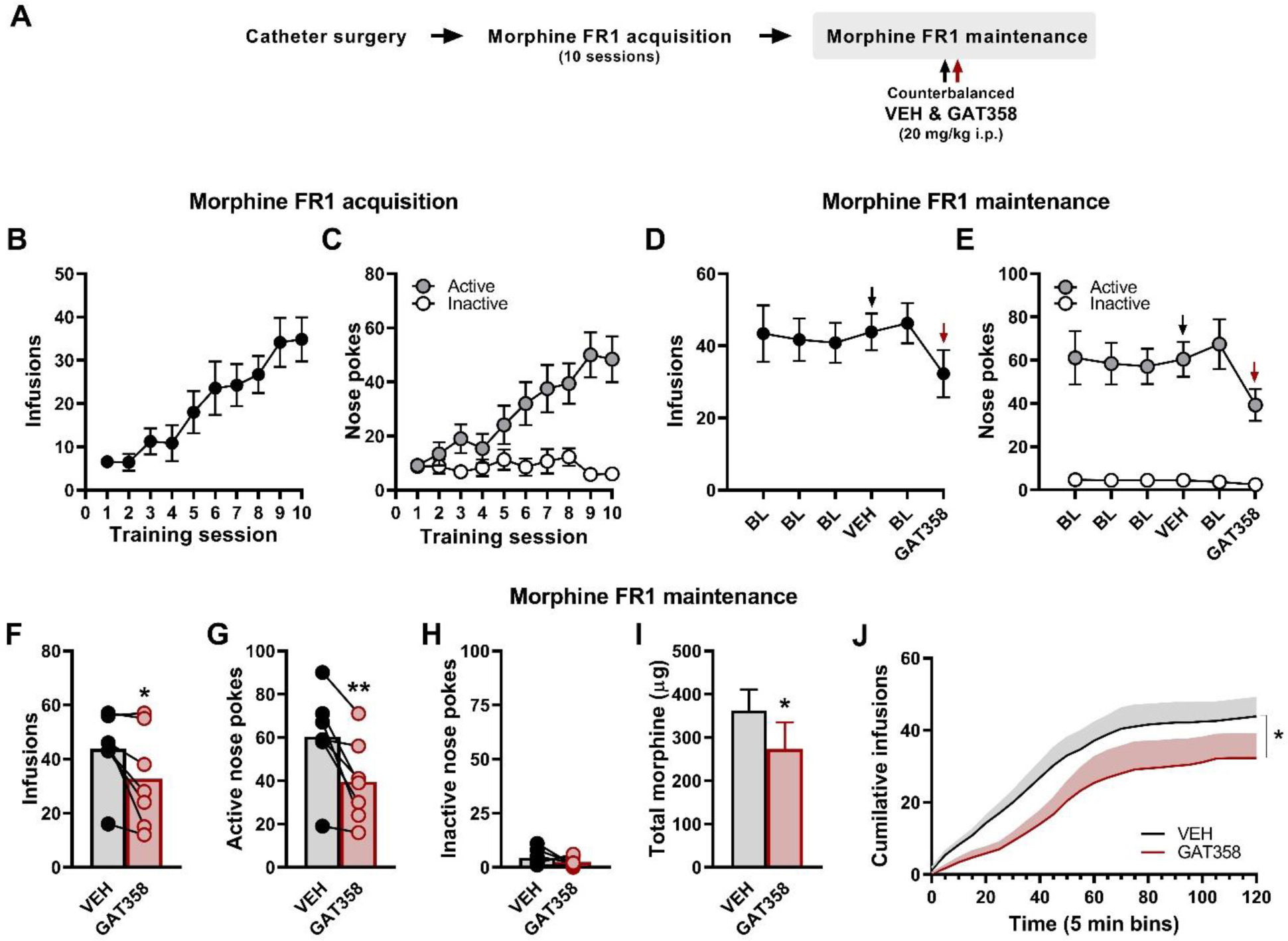
GAT358 reduces morphine consumption after the establishment of stable drug infusion intake. **(A)** Schematic shows the order of the experimental procedures. Mice were implanted with intravenous jugular catheters. After recovering from the surgery, mice were allowed to self-administer morphine under FR1 for 10 sessions (acquisition phase). During the maintenance phase, mice were allowed to continue self-administering morphine under FR1 until reaching a stable behavior (>10 infusions and less than or equal to 15% variance for 3 consecutive days). Subsequently, all animals received GAT358 (20 mg/kg i.p.) and vehicle (VEH, i.p.) in a counterbalanced manner 20 min before behavioral sessions. Number of morphine infusions (**B**) and number of active and inactive nose pokes (**C**) across the initial 10 morphine self-administration sessions comprising the acquisition phase. Number of morphine infusions (**D**) and number of active and inactive nose pokes (**E**) during the maintenance phase of morphine self-administration after reaching stable baseline (**BL**) responding and after receiving counterbalanced injections of vehicle (VEH, i.p.) and GAT358 (20 mg/kg i.p.). Note that in the figure, data are normalized to show vehicle (VEH) first, but treatments were counterbalanced between animals. Average number of morphine infusions (**F**), average number of active nose pokes (**G**) and average number of inactive nose pokes (**H**) during the maintenance phase of self-administration in mice treated with GAT358 (20 mg/kg i.p.) and vehicle (VEH, i.p.). **(I)** Total morphine consumed in the maintenance phase of self-administration after the treatment with GAT358 (20 mg/kg i.p.) and vehicle (VEH, i.p.). **(J)** Average cumulative infusions of animals that received GAT358 (20 mg/kg i.p.) vs. vehicle (VEH, i.p.) in the maintenance phase of morphine self-administration. Data are expressed as mean ± SEM (n = 7). *p <0.05, **p <0.01, ***p <0.001.

### 3.2. GAT358 decreases the relapse of morphine seeking after forced abstinence

Next, we studied the impact of GAT358 (20 mg/kg i.p.) on the relapse of morphine seeking after 21 days of forced abstinence (**Figure 2**). In this experiment, animals self-administered morphine for 15 days under FR1, were subjected to 21 days of forced abstinence in their home cage and then injected with GAT358 (20 mg/kg i.p.) or vehicle (i.p.) prior to undergoing a single relapse test session (**Figure 2A**). After the initial 15 sessions of drug self-administration, animals successfully acquired morphine intake behavior (**Figure 2B**) and accurately discerned the reinforced from the non-reinforced nose poke throughout the 15 self-administration sessions (**Figure 2C**). Pre-treatment with GAT358 (20 mg/kg i.p.) successfully decreased morphine-seeking behaviors in a relapse test (**Figure 2D**). A two-way ANOVA revealed main effects of treatment (F_1,40_=13.53, p=0.0007) and nose poke type (F_1,40_=34.84, p<0.0001) and a significant interaction between treatment and the nose poke type (F_1,40_=6.369, p=0.0157) were also observed. Bonferroni’s post-hoc multiple comparisons revealed that the number of active nose pokes (p=0.0002) was lower in the GAT358 (20 mg/kg i.p.)-treated compared to the vehicle-treated group. By contrast, the number of inactive nose pokes did not differ between groups (p=0.8383) (**Figure 2D**).

**Figure 2.**
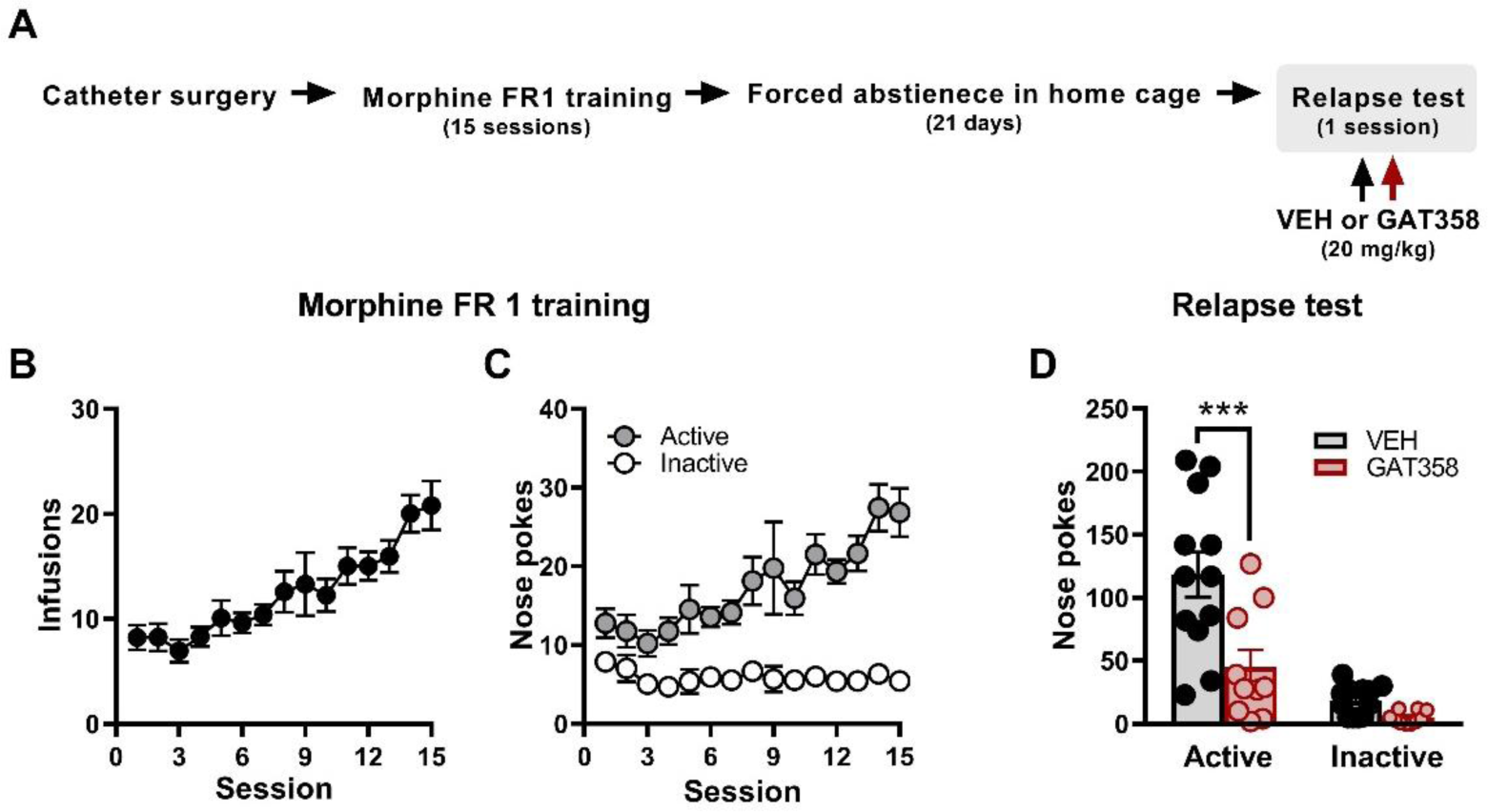
GAT358 decreases the relapse of morphine seeking behavior after forced abstinence. **(A)** Schematic shows the order of experimental procedures. Mice were implanted with intravenous jugular catheters. After recovering from the surgery, animals were allowed to self-administer morphine under FR1 for 15 session and then spent 21 days in forced abstinence in their home cages. Then, mice were subjected to a single relapse session in which they received vehicle (VEH) or GAT358 (20 mg/kg i.p.) 20 min before behavioral testing. Morphine infusions intake (**B**) and active and inactive nose pokes (**C**) during FR1 training. **(D)** Number of active and inactive nose pokes during the relapse test in animals treated with vehicle (VEH) or GAT358 (20 mg/kg i.p.). Data are expressed as mean ± SEM, VEH n=12, GAT358 n=10. *p <0.05, **p <0.01, ***p <0.001.

### 3.3. GAT358 reduced the motivation to obtain morphine rewards

We evaluated the effects of different doses of GAT358 on the motivation to work for morphine rewards (**Figure 3**). In this experiment, mice were first trained to self-administer morphine under FR1 (10 days), FR3 (5 days) and FR3 (5 days), and then subjected to PR, where they received counterbalanced doses of GAT358 (10, 20 and 30 mg/kg i.p.) (**Figure 3A**). At the end of the FR training, mice increased the number of infusions taken (**Figure 3B**) and clearly recognized the difference between active and active nose pokes (**Figure 3C**). One-way ANOVA, comparing the last two sessions in each FR, showed no differences in the number of infusions between FRs (F_2,18_=0.3042, p=0.7414; **Figure 3D**). However, the number of active nose pokes was higher (F_2,18_=4.45, p=0.0269) under FR3 compared to FR1 schedules (p=0.0239; **Figure 3E**). The number of inactive nose pokes (F_2,18_=4.483, p=0.0263) was similarly lower under FR3 compared to FR1 schedules (p=0.0234; **Figure 3F**). In the PR schedule, GAT358 (10, 20, 30 mg/kg i.p.) dose-dependently decreased the number of morphine infusions taken (**Figure 3G**). GAT358 treatment also decreased the number of morphine infusions (F_1.89,11.38_= 17.68, p=0.0004); GAT358 30 mg/kg i.p. reduced infusion intake compared to either vehicle (p=0.0004), GAT358 10 mg/kg i.p. (p=0.0088) or GAT358 20 mg/kg i.p. (p=0.0145). Furthermore, we observed a significant negative linear trend between treatments (p<0.0001; **Figure 3G**). Similarly, GAT358 (10, 20, 30 mg/kg i.p.) dose-dependently decreased the number of active nose pokes (F_1.75,10.51_=6.487, p=0.0168; **Figure 3H**); GAT358 30 mg/kg i.p. decreased the number of active nose pokes compared to vehicle (p= 0.0238), GAT358 10 mg/kg i.p. (p= .0088) and GAT358 20 mg/kg i.p. (p= 0.0464). A negative linear trend was also observed between treatments (p<0.0004; **Figure 3H**). By contrast, the number of inactive nose pokes remained almost unaltered across the different doses of GAT358 (F_1.52,9.16_=4.37, p=0.0540; **Figure 3I**). GAT358 also decreased the break point (i.e., final ratio completed) in a dose-dependent manner (F_1.51,9.07_=10.36, p=0.0064; **Figure 3J**). The breakpoint was lower in GAT358 30 mg/kg i.p. treated groups compared to vehicle (p= 0.0079), GAT358 10 mg/kg i.p. (p=0.0356) and GAT358 20 mg/kg i.p. (p=0.0113) treated groups. A negative linear trend in breakpoint was also observed between treatments (p<0.0001; **Figure 3J**).

**Figure 3.**
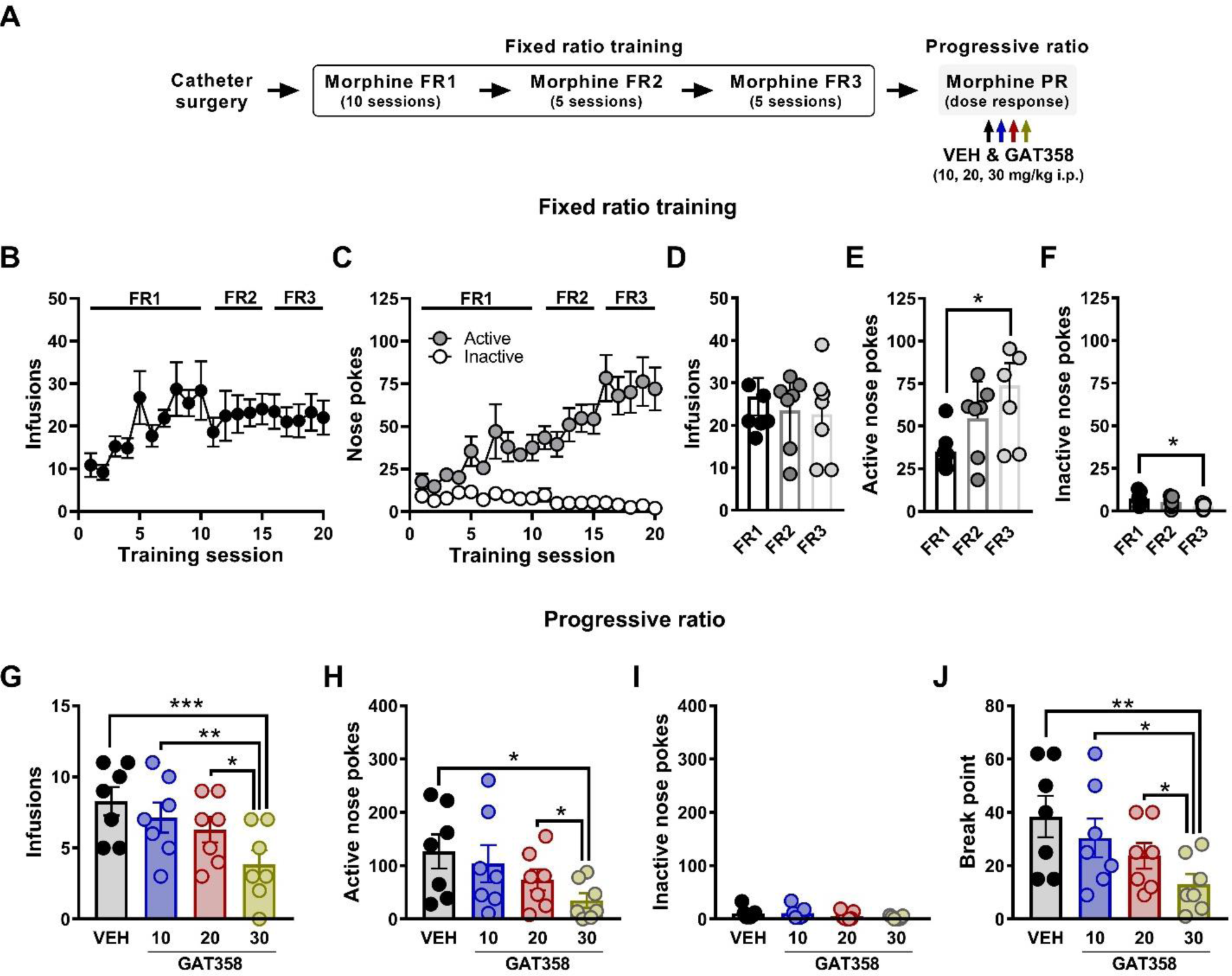
GAT358 reduces the motivation to work for morphine rewards in a progressive ratio schedule of reinforcement. **(A)** Schematic shows the experimental design. Briefly, mice were first trained under FR1 (10 days), FR2 (days) and FR3 (5 days). Then mice underwent to PR training until reaching stable morphine infusion intake (i.e., less than 15% variance for 3 consecutive days). After mice acquired stable behavior under PR, they received counterbalanced injections of VEH (i.p.) and GAT358 (10, 20 and 30 mg/kg i.p.) before behavioral testing. **(B)** Infusion intake increased during the fixed ratio training under FR1 and was stable under FR2 and FR3. **(C)** Active nose pokes were greater than inactive nose pokes during the fixed ratio training. Average number of infusions (**D**), active nose pokes (**E**), and inactive nose poke in the last 2 days of each fixed ratio training (**F**). Effect of GAT358 doses on morphine infusions earned (**G**), the number of active nose pokes (**H**), the rate of inactive nose pokes (**I**) and breakpoint for morphine infusion intake (**J**) during PR schedule of reinforcement. Data are expressed as mean ± SEM (n = 7). *p <0.05, **p <0.01, ***p <0.001.

### 4. GAT358 did not produce motor ataxia

GAT358 (10, 20, and 30 mg/kg i.p.) did not impair motor coordination in the rotarod test (**Figure 4A**). No differences were detected in latency to descend from the rotating drum in groups treated with either vehicle or any dose of GAT358 (10, 20, and 30 mg/kg i.p.) (**Figure 4A, B**). Two-way ANOVA did not detect an interaction in descent between pre- and post-injection trials across treatments (F_3,27_=0.9577, p=0.4269; **Figure 4, B**).

**Figure 4.**
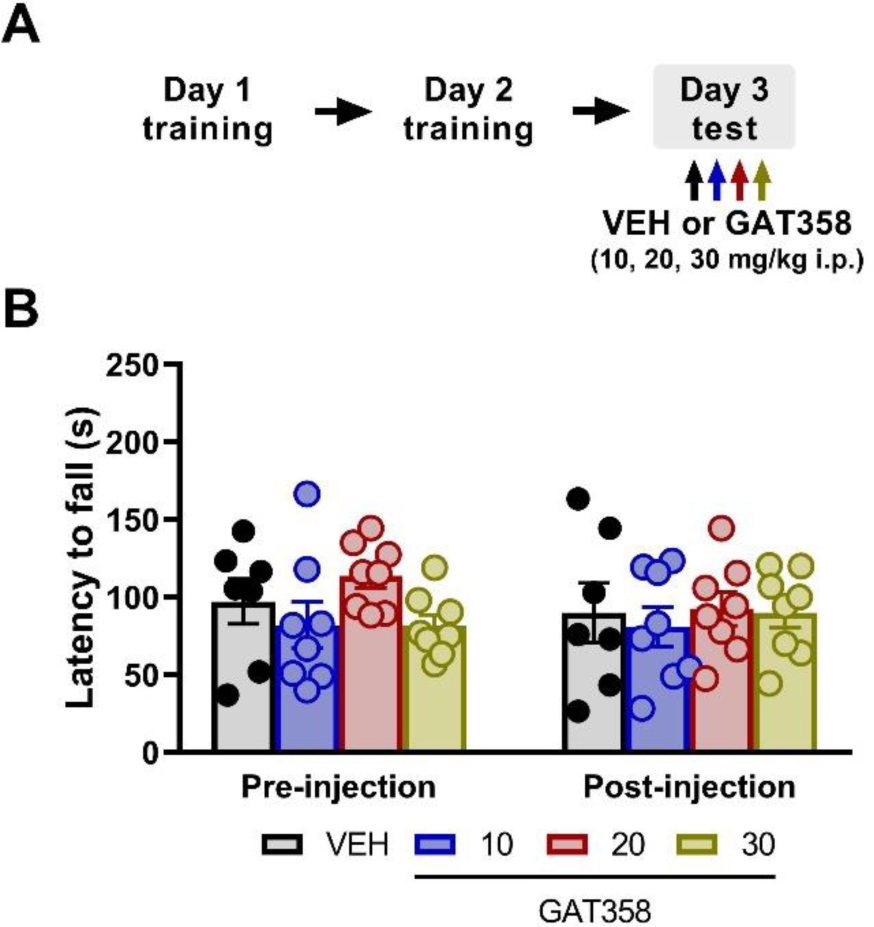
GAT358 did not produce motor ataxia in the rota-rod test. **(A)** Schematic shows the experimental timeline. **(B)** The latency to descend from the drum did not differ between GAT358 (10, 20 or 30 mg/kg i.p.) or vehicle treatment during either pre-injection or post-injection trials. Data are expressed as mean ± SEM, VEH n=7, GAT358 10mg/kg n=8, GAT358 20mg/kg n=8, GAT358 30mg/kg n=8.

### 5. GAT358 did not reduce the motivation to work for food rewards

We asked whether GAT358 would alter the self-administration of food rewards (**Figure 5A**). In this experiment, mice underwent training to work for food (i.e., sucrose pellets) under FR1, FR2, and FR3 schedules, then transitioned to a PR schedule until achieving stable pellet intake when they received VEH i.p., or GAT358 (10, 20, and 30 mg/kg i.p.) (**Figure 5A**). None of the doses of GAT358 altered the number of pellets consumed (F_1.9,11.47_=0.7913, p=0.4712; **Figure 5B**), the number of active nose pokes (F_1.9,11.65_=0.3819, p=0.6850; **Figure 5C**), the number of the inactive nose pokes (F_1.1,6.8_=0.4931, p=0.594; **Figure 5D**), or the motivation to get food rewards (F_1.8,11.2_=0.4463, p=0.6349; **Figure 5E**).

**Figure 5.**
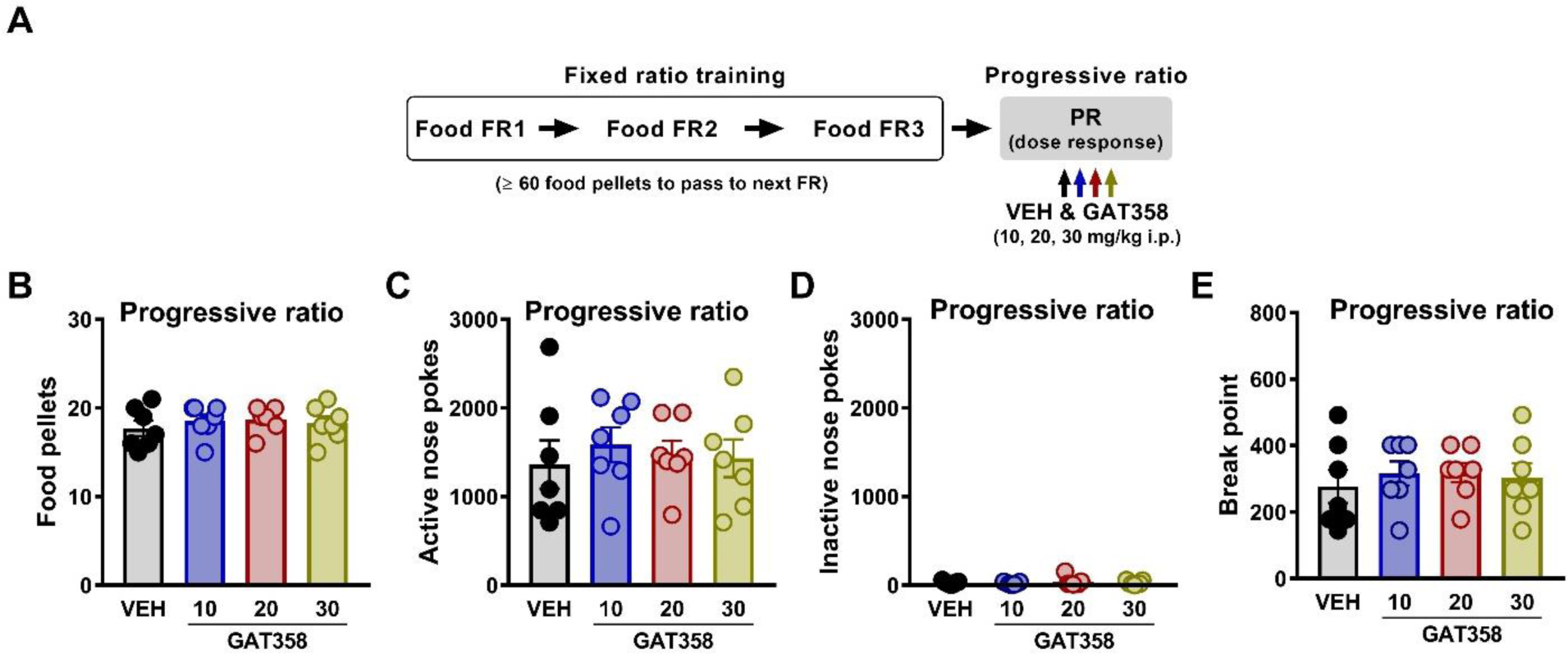
GAT358 did not affect the motivation to obtain food rewards in a progressive ratio schedule of reinforcement. **(A)** Schematic shows the sequence of the experimental procedures. Mice were first trained under FR1, FR2 and FR3 with the requisite to get 60 or more food pellets to pass from one FR to the next. Then mice underwent to PR training until they reached stable food pellet intake. After acquiring stable behavior under PR, mice were treated in a counterbalanced manner with VEH (i.p.) and GAT358 (10, 20, and 30 mg/kg i.p.) 20 min prior to behavioral testing. GAT358 did not alter the number of sucrose pellets earned **(B)**, the number of active nose pokes (C), the number of inactive nose pokes **(D)** or the break point **(E)** during PR schedule of reinforcement at any dose. Data are expressed as mean ± SEM, n=7 for all groups.

## 4. Discussion

CB1 NAMs have been included among the medication development priorities of the National Institute on Drug Abuse (NIDA) in response to the opioid crisis ^18^. CB1 NAMs represent a potential alternative or adjunct treatment to opioids, with the potential to produce a positive impact on society by furthering efforts to combat the opioid epidemic. It is widely recognized that OUD disrupts brain-motivated behaviors and impairs the internal control that a person has over the substance, effects presumably caused by changes in the reinforcing efficacy of the abused drug ^19^. In the present study, we examined the impact of the CB1-biased NAM GAT358 on the reinforcing properties of morphine using a self-administration operant task in which mice received i.v. morphine infusions upon completing a specific number of nose pokes. The self-administration paradigm allowed us to evaluate the effects of GAT358 in several aspects of morphine addiction, such as drug intake, relapse, or motivational strength. The results presented in this study support the hypothesis that the CB1 NAM GAT358 may be leveraged as an effective treatment to decrease morphine addictive behaviors.

GAT358, when administered (i.p.) during the maintenance phase of drug self-administration (i.e., when there is an ongoing stable morphine intake), reduced the number of morphine infusions taken, the number of active nose pokes and the total morphine consumed compared to vehicle. Furthermore, GAT358 reduced the escalation of morphine infusion intake and the maximum number of cumulative morphine infusions received. Moreover, when we increased the schedule of reinforcement using a progressive ratio task, GAT358 dose-dependently decreased the number of infusions taken, the number of active nose pokes and the motivation (i.e., break point) to work to get morphine infusions. These results are consistent with our previous studies that demonstrated that GAT358 decreased oxycodone consumption in a two-bottle choice, prevented CPP to morphine and eliminated morphine-induced increases in electrically-evoked dopamine efflux in the mesocorticolimbic pathway ^14^. To our knowledge this is the first time that the effects of a CB1R NAM on the motivation to obtain opioids have been evaluated. Importantly, our data supports the hypothesis that therapeutic interventions based on CB1R NAMs may reduce opioid reinforcement and motivation to self-administer opioids in the context of morphine addiction.

The ability of GAT358 to reduce opioid-addicted behaviors was also maintained when morphine was withheld during the relapse test. GAT358 decreased the actions in the previously morphine-paired nose poke, and ultimately diminished morphine-seeking behavior to drug-cue presentations during abstinence. As far as we are aware, this is the only report to examine the actions of a CB1R NAM on relapse to opioids. Our findings align with previous studies demonstrating a reduction in the reinstatement of cocaine- and methamphetamine-seeking behavior in rats employing other CB1R NAMs, such as PSNCBAM-1 and Org27569 ^6,7^.

Other CB1R NAMs, including PSNCBAM-1 or Org27569, have been associated with decreases in both food intake and locomotor activity, side effects that are believed to be caused by its inverse agonist profile ^20^. However, it is unlikely that any of the outcomes described here with GAT358 result from nonspecific motor impairment or other adverse psychological effects. GAT358 showed a minimal CB1 inverse agonist profile and displayed functional selectivity in the β-arrestin assay ^13^. In our study, GAT358 did not reduce the number of inactive nose pokes in the morphine maintenance, relapse, or progressive ratio experiments, which, to some extent, lessens the possibility of nonspecific motor impairment. Nonetheless, we further assessed motor function using the rota-rod test since motor ataxia hinders the execution of tasks demanding precision and coordination, like those faced in an operant task paradigm. None of the GAT358 doses employed here altered the latency to descend from the rotarod and did not produce motor ataxia in any instance. Furthermore, we confirmed that none of the doses of GAT358 evaluated herein reduced the motivation to work for food rewards, or decreased the number of pellets consumed, consistent with absence of drug-induced anhedonia. Together, these findings suggest that GAT358 may have a more beneficial pharmacological profile over other existing CB1R NAMs. More research is necessary to elucidate if the effects of GAT358 described here are dependent upon the β-arrestin-biased pharmacological profile of this ligand.

GAT358 not only reduced opioid-addicted behaviors when morphine was present and self-administered, but also when morphine was withheld during the relapse test. As such, this observation suggests that negative allosteric modulation of CB1 cannabinoid receptor signaling influences distinct brain pathways involved in reward processing and seeking. In the VTA, endocannabinoids retrogradely modulate both glutamatergic excitatory and GABAergic inhibitory synaptic inputs into dopamine neurons ^5^. The actions of GAT358 on the VTA may account for the primary rewarding effects observed in this study. In the NAc, endocannabinoids retrogradely modulate the glutamatergic afferents from prefrontal regions onto D1-medium spiny neurons in a dopamine-independent mechanism ^21^. Increased glutamate levels in the NAc have been associated with cue-induced reinstatement and drug-seeking behaviors ^22–24^. Hence, GAT358 could be modulating glutamatergic afferents onto the NAc during the relapse test. This hypothesis is consistent with the selective effects of disruption of protein-protein interactions downstream of NMDARs that suppress aberrant glutamate excitability on relapse of morphine-seeking behavior ^25^. CB1Rare also present in the prefrontal cortex, which processes the hedonic value and motivation to obtain drugs ^26^. Thus, GAT358 could be acting in the prefrontal cortex to decrease the motivation to work for drug rewards. Given that GAT358 was administered systemically in our experiments, either direct or indirect mechanisms could mediate its effects on the reinforcing properties of morphine described in this study. Likewise, our results do not preclude the possibility that GAT358 may differentially impact distinct neuronal populations to mediate the effects observed herein. Further research is warranted to elucidate the specific cell types impacted by GAT358 and to characterize the underlying circuit mechanisms. Future studies are also essential to determine if the results observed in male mice generalize to female mice.

In addition to the results presented here, a recent study from our lab also explored the application of the CB1R NAM GAT358 on antinociceptive and effects of morphine as well as morphine tolerance and naloxone-precipitated opioid withdrawal ^27^. We showed that GAT358 did not impede morphine antinociception but was effective in reducing morphine tolerance as well as naloxone-precipitated opioid withdrawal ^27^. As such, GAT358 provide new options for non-addictive pain management, as it produced antinociception on its own, and also potentially reduce the reliance and addiction liability of opioids. These observations raise the possibility that combination treatments incorporating CB1R NAMs and opioids have the potential to lower the risk of addiction liability, while exhibiting therapeutic potential for more effective pain reduction.

## 5. Conclusion

Collectively, our findings demonstrate that GAT358 effectively decreased morphine self-administration when there is ongoing stable morphine consumption, reduced morphine relapse-like behavior, and lowered the motivation to work to obtain morphine infusions. Importantly, GAT358 did not produce side effects like motor ataxia or anhedonia that could confound our results. This study highlights that novel therapies based on CB1R NAMs have the potential to offer a favorable safety profile and are viable for reducing the risk associated with opioid use.

## Abbreviations

CB1: cannabinoid-type 1 receptor
NAM: negative allosteric modulator
SUD: substance use disorder
OUD: opioid use disorder
FR: fixed ratio
PR: progressive ratio

## Acknowledgements

The authors are grateful to Emily Fender and John Hainline for providing technical assistance. Graphical abstract was created with Biorender.com.

## Credit authorship contribution statement

IO performed drug/food self-administration and rota-rod experiments with assistance from FK. LNC and GAT designed and synthesized GAT358. IO and AGH designed the experiments. IO and JDC wrote programs for data collection and analysis for the drug/food self-administration studies. AGH supervised the project. IO and AGH wrote the manuscript with assistance from JDC and GAT.

